# Age-dependent increase in α-tocopherol and phytosterols in maize leaves exposed to elevated ozone pollution

**DOI:** 10.1101/2020.11.09.360644

**Authors:** Jessica M. Wedow, Charles H. Burroughs, Lorena Rios Acosta, Andrew D.B. Leakey, Elizabeth A. Ainsworth

## Abstract

Tropospheric ozone is a major air pollutant that significantly damages crop production around the world. Crop metabolic responses to rising chronic ozone stress have not been well-studied in the field, especially in C_4_ crops. In this study, we investigated the metabolomic profile of leaves from two diverse maize (*Zea mays*) inbred lines and the hybrid cross during exposure to season-long elevated ozone (~100 nL L^−1^) in the field using free air concentration enrichment (FACE) to identify key biochemical responses of maize to elevated ozone. Senescence, measured by loss of chlorophyll content, was accelerated in the hybrid line, B73 x Mo17, but not in either inbred line (B73 or Mo17). Untargeted metabolomic profiling further revealed that inbred and hybrid lines of maize differed in metabolic responses to ozone. A significant difference in the metabolite profile of hybrid leaves exposed to elevated ozone occurred as leaves aged, but no age-dependent difference in leaf metabolite profiles between ozone conditions was measured in the inbred lines. Phytosterols and α-tocopherol levels increased in B73 x Mo17 leaves as they aged, and to a significantly greater degree in elevated ozone stress. These metabolites are involved in membrane stabilization and chloroplast reactive oxygen species (ROS) quenching. The hybrid line also showed significant yield loss at elevated ozone, which the inbred lines did not. This suggests that the hybrid maize line was more sensitive to ozone exposure than the inbred lines, and up-regulated metabolic pathways to stabilize membranes and quench ROS in response to chronic ozone stress.

## Introduction

Tropospheric ozone (O_3_) is a secondary air pollutant formed from the reaction of nitrogen oxides and volatile organic compounds in the presence of UV radiation. From 1980 to 2011, O_3_ concentrations ([O_3_]) in the United States are estimated to have caused a yield loss of up to 10% for rain-fed maize, with an estimated economic cost of $7.2 billion per year (McGrath *et al.*, 2015). While much is known about the metabolic and signaling responses within plant cells to elevated [O_3_], major knowledge gaps remain with regard to: (1) the mechanisms underlying genetic variation in O_3_ response, and (2) the nature of biochemical O_3_ responses in the production environment of a farm field (Ainsworth, 2017). Additionally, our understanding of plant metabolic responses to O_3_ stress largely comes from acute experiments in controlled environments (Vainonen and Kangasjärvi, 2015), yet it is well known that the mechanisms of response to chronic O_3_ exposure, generally defined as long-term exposure to concentrations of ~100 nL L^−1^ or less, differ from acute responses to very high [O_3_] (Vahala *et al.*, 2003; Grantz and Vu, 2012). Increased emissions of precursor pollutants have primarily increased crop exposure to chronic O_3_ stress, with tropospheric [O_3_] increasing from ~10 nL L^−1^ in the late 1800’s to ~40-50 nL L^−1^ today (Monks *et al.*, 2015; Brauer *et al.*, 2016). In many countries, [O_3_] continue to increase (Brauer *et al*., 2016), further intensifying chronic O_3_ exposure.

Ozone diffuses through the stomata into the intercellular airspace where it rapidly reacts to form additional reactive oxygen species (ROS). It is also a powerful oxidizing agent capable of reacting with diverse molecules including lipids, proteins, nucleic acids and carbohydrates. ROS formed from O_3_ further react with apoplastic antioxidants and a number of proteins embedded in the plasma membrane (e.g., NADPH oxidases, aquaporins, receptor-like kinases, and G-proteins) and elicit an increase in cytosolic calcium (Short *et al.*, 2012). Peroxidation and denaturation of membrane lipids can also occur with prolonged or acute O_3_ exposure (Pell *et al.*, 1997; Loreto and Velikova, 2001). The intracellular response to the influx of ROS depends upon the duration and intensity of O_3_ exposure, with calcium, hormone signaling, and MAP kinase cascades all playing a role in regulation of transcriptional and biochemical changes (Vainonen and Kangasjärvi, 2015). Under chronic O_3_ stress, transcriptional changes have been associated with decreased photosynthesis (Pell *et al.*, 1997; Leitao *et al.*, 2007; Li *et al.*, 2019), increased rates of mitochondrial respiration and antioxidant production (Gillespie *et al.*, 2012; Yendrek *et al.*, 2015), increased hormone biosynthesis (jasmonates, ethylene, and salicylic acid) (Kangasjärvi *et al.*, 2005; Vainonen and Kangasjärvi, 2015), and early activation of several *senescence associated genes* (SAGs) (Miller *et al.*, 1999; Fiscus *et al.*, 2005; Kangasjärvi *et al.*, 2005; Betzelberger *et al.*, 2012; Gillespie *et al.*, 2012). Shifts in metabolism from carbon assimilation to defense and detoxification in combination with early senescence, are thought to be major drivers of reduced plant productivity under elevated [O_3_] (Dizengremel, 2001; Morgan *et al.*, 2006; Feng *et al.*, 2011; Yendrek *et al.*, 2017a; Choquette *et al*., 2019; Choquette *et al*., 2020).

A wide range of metabolites have been reported to change in response to elevated [O_3_] in different species, many in common with plant defense responses (Iriti and Faoro, 2009). Plant steroids, including phytosterols and brassinosteroids (BRs), increase plant tolerance to a wide range of abiotic and biotic stresses as well as control plant growth, flowering time and senescence (Vriet *et al.*, 2012). Changes in the ratios of membrane steroids during stress is also commonly reported following abiotic and biotic stress treatments (Rogowska and Szakiel, 2020). For example, the ratio of stigmasterol to β-sitosterol increased following exposure of Arabidopsis to pathogen-associated molecular patterns and ROS, which was thought to help maintain plasma membrane fluidity and permeability during stress (Griebel and Zeier, 2010). In the chloroplasts, α-tocopherol is an important scavenger that protects photosynthetic machinery by quenching singlet oxygen or inhibiting the progression of lipid peroxidation (Havaux *et al.*, 2005). Exposure to many abiotic stresses that increase ROS results in increased α-tocopherol content in leaves (Munné-Bosch, 2005). As leaves age, α-tocopherol content also increases (García-Plazaola and Becerril, 2001; Hormaetxe *et al*., 2005). It is unknown if these metabolites play a role in O_3_ response under chronic exposure and field conditions. If so, breeding or biotechnology might be used to leverage the protective roles of phytosterols and non-enzymatic antioxidants to improve tolerance to O_3_-induced oxidative stress.

More broadly, metabolomics provides a tool to explore biochemical signatures that may be predictive of environmental stress effects on primary productivity, even in field conditions. Studies have examined the relationship between leaf metabolites and physiological traits in the field under various abiotic stress conditions in maize (Riedelsheimer *et al.*, 2012; Obata *et al.*, 2015), rice (Melandri *et al.*, 2020), and Guinea grass (Wedow *et al.*, 2019). But, metabolomic responses of maize to elevated [O_3_] have not yet been widely characterized despite that maize is one of the world’s most widely-grown crops (USDA FAS, 2020). Maize is also a model species for the C_4_ plant functional group that includes many other important crops used to produce food, fuel, forage and fiber. Therefore, we examined the metabolomic profile of maize inbred (B73 and Mo17) and hybrid (B73 x Mo17) lines grown at a Free Air Concentration Enrichment (FACE) facility at the University of Illinois, Urbana-Champaign. The elevated [O_3_] treatment was roughly 2.5 times the current average summer concentration in central Illinois, and consistent with current [O_3_] in polluted regions of Asia (Hong *et al.*, 2019). Additionally, many regions that currently experience high O_3_ events overlap with the most productive croplands across the world (Ramankutty *et al.*, 2008; Ainsworth, 2017).

In this study, leaf metabolomic profiles were investigated at three time points from early to mid-stages of leaf senescence as characterized by chlorophyll content. Metabolite content was then linked to leaf mass per unit area and grain yield to identify potential biochemical markers for O_3_ response in maize. B73 and Mo17 are two classic elite inbred maize lines (Stuber *et al.*, 1992), which serve as the parents for widely used mapping populations to study the genetic architecture of numerous traits (Mickelson *et al.*, 2002; Balint-Kurti *et al.*, 2007; Wassom *et al.*, 2008; Pressoir *et al.*, 2009; Sorgini *et al.*, 2019). We aimed to test the hypotheses that (1) maize metabolomic signatures would be altered under elevated [O_3_], with more pronounced effects as the leaves aged; (2) an increase in metabolites associated with oxidative stress and/or membrane stabilization would occur in sensitive maize lines; (3) key metabolites would be correlated with variation in yield loss under elevated [O_3_] providing potential molecular markers of O_3_ response in maize.

## Materials and methods

### Field site and experimental conditions

In 2015, maize (*Zea mays*) inbred (B73 and Mo17) and hybrid (B73 x Mo17) lines were studied at the FACE research facility in Savoy, IL (https://soyface.illinois.edu/). The facility is a 32-ha farm where maize and soybean (*Glycine max*) are grown in annual rotation. Maize inbred and hybrid lines were planted with a precision planter in rows spaced 0.76 m apart and 3.35 m in length at a density of 8 plants m^−1^. There were four pairs of ambient [O_3_] (~40 nL L^−1^) and elevated [O_3_] (100 nL L^−1^) plots (*n* = 4) in the experiment (Figure S1). Inbred and hybrid lines were grown in separate plots to avoid the taller hybrids altering the light environment or interfering with fumigation of shorter inbred lines, which resulted in the use of 16 octagonal, 20 m diameter plots. One replicate pair of ambient and elevated [O_3_] plots with the hybrid line was dropped from the analysis due to water logging. Each genotype was planted in five different locations within each plot (Figure S1). Additional site information and field conditions were described in Yendrek *et al*. (2017a).

Air enriched with O_3_ was delivered to the experimental rings with FACE technology as described in Yendrek *et al.* (2017b). The target elevated [O_3_] was 100 nL L^−1^ and the O_3_ treatment was administered from 10:00 to 18:00 throughout the growing season when it was not raining and the wind speed was greater than 0.5 m s^−1^. Based on 1 min average [O_3_] collected in each ring, the fumigation was within 10% of the 100 nL L^−1^ target for 56% of the time and within 20% of the target for 79% of the time in 2015. Weather conditions were measured with an on-site weather station as reported previously in Yendrek *et al*. (2017a).

### Sampling protocol and tissue handling

Leaf material for metabolomic profiling of B73, Mo17 and B73 x Mo17 was sampled on three dates corresponding to leaf physiological maturity, early- and mid-senescence. Samples were taken in the inbred experiment on: 4 August 2015, 13 August 2015, and 24 August 2015 (day of year DOY: 216, 225, and 236) and in the hybrid experiment on: 27 July 2015, 7 August 2015, and 17 August 2015 (DOY): 208, 219, and 229). Different plants within the replicate rows were sampled on each date. All material was collected from the plots between 12:00 and 14:00. Approximately 100 cm^2^ of leaf material was cut from the middle of the leaf subtending the ear with scissors, wrapped in tinfoil and immediately placed in liquid nitrogen, before being stored at −80°C. Five samples per inbred and hybrid line (1 per sub-block) were taken from each ambient and elevated [O_3_] plot.

Leaf mass per unit area (LMA) was measured from leaf disks taken at the same time as samples for metabolomic profiling. Three leaf disks (0.02 m dia) per row were cut with cork borers, placed into coin envelopes and dried in an oven at ~60°C for one week. Samples were then weighed and data from the 5 rows within each plot were averaged for a plot-level estimation of LMA (g m^−2^).

### Leaf chlorophyll content

Chlorophyll content was estimated from measurements collected with a SPAD meter (Konica-Minolta SPAD-502 Chlorophyll meter, Japan). Measurements were collected from the leaf subtending the ear of three plants per genotype at five locations per ring. Six readings were taken from the middle third of each leaf and the average value was recorded. Measurements were taken approximately every three days starting at anthesis, continuing through leaf senescence (DOY 204-255). The equation: chlorophyll content (μg cm^−2^) = (99*SPAD)/(144-SPAD) was used to convert SPAD values to chlorophyll content (Cerovic *et al.*, 2012).

### Seed yield

At maturity, ears were harvested from 8 plants in each of the 5 rows per genotype per plot, dried for ~1 week in an oven at ~60°C, then shelled and weighed to estimate yield (g plant^−1^).

### Metabolomic profiling by GC-MS

Untargeted metabolomic profiling was performed with 15 mg of lyophilized leaf tissue according to the protocol described in Ulanov and Widholm (2010). A total of 40 samples per time point for each inbred, B73 and Mo17, and 30 samples per time point for B73 x Mo17 were processed for metabolomic profiling. Samples were extracted with methanol: acetone: H_2_O (1:2:1, v/v/v) at ambient temperature. For quality control (QC), 10 μl of leaf extract was taken from each sample and pooled, then run and analyzed after every 9 biological samples. Samples and QC were dried under vacuum and derivatized with 75 μl methoxyamine hydrochloride (Sigma-Aldrich, MO, USA) (40 mg ml^−1^ in pyridine) for 90 min at 50°C, then with 125 μl MSTFA + 1%TMCS (Thermo, MA, USA) at 50°C for 120 min followed by an additional 2-h incubation at room temperature. An internal standard (30 μL hentriacontanoic acid) was added to each sample prior to derivatization. Samples were analyzed on a gas chromatography/mass spectroscopy (GC/MS) system (Agilent Inc, Palo Alto, CA, USA) consisting of an Agilent 7890 gas chromatograph, an Agilent 5975 mass selective detector, and a HP 7683B autosampler. Gas chromatography was performed on a ZB-5MS capillary column (Phenomenex, Torrance, CA, USA). The inlet and MS interface temperatures were 250 °C, and the ion source temperature was adjusted to 230 °C. An aliquot of 1 μl was injected with the split ratio of 10:1. The helium carrier gas constant flow rate was 2.4 ml min^−1^. The temperature program was 5 min isothermal heating at 70 °C, followed by an oven temperature increase of 50°C min^−1^ to 310 °C, and a final 10 min at 310 °C. The mass spectrometer was operated in a positive electron impact mode at 69.9 eV ionization energy in m/z 50-800 scan range.

Raw data files were processed with the metaMS.GC workflow hosted on the workflow4metabolomics (W4M) server (Giacomoni *et al.*, 2015). Default settings were used except for minimum class fraction, specified at 0.6. Spectra were normalized to the internal standard and leaf dry weight was used to account for tissue and water content differences over time. Batch correction was done with the all loess sample regression model, W4M tool. Peak annotation used a custom-built database and AMDIS 2.71 (NIST, Gaithersburg, MD, USA) program. All known artificial peaks were identified and removed. The instrument variability was within the standard acceptance limit of 5%.

### Statistical analysis

The analysis of chlorophyll loss over time was tested by fitting a quadratic equation to the data where: Chl = y_0_ + α*x + β*x^2^, where x equals day of year. To test for differences in chlorophyll loss over time in ambient and elevated [O_3_], a single quadratic model was first fit to the data for each genotype (PROC NLIN, SAS 9.4, SAS Institute, Cary, NC), then models were fit to each genotype and treatment combination. An F statistic was used to test if the model with genotype and treatment produced a significantly better fit to the data, that is, if there were significant differences in the response of chlorophyll over time in ambient and elevated [O_3_], following the approach of Yendrek *et al*. (2017a).

LMA and seed yield were analyzed using analysis of variance. For the inbred experiment, the model included fixed effect terms for inbred line and treatment, and a random term for block. The model for the hybrid experiment included treatment as a fixed effect and block as a random term in the model. Significant differences between treatments were determined by Tukey tests with a threshold of p<0.05.

The inbred and hybrid metabolite data were analyzed separately because they were grown in different field plots and harvested on different dates (Figure S1). For each dataset, metabolite data were log_10_ transformed and processed with univariate analysis to identify and remove outliers (studentized residual ≥ 4). The 5 observations for a genotype within each FACE or control plot were averaged for analysis and time points were analyzed separately. Each metabolite was tested independently using a two-way ANOVA (Kirpich *et al.*, 2018) for the inbred experiment with treatment and genotype as fixed effects, and a one-way analysis of variance model for the hybrid experiment (Figure S2). Statistical differences in least squared mean estimates between ambient and elevated [O_3_] for each time point were determined by Tukey tests with a threshold of *p* < 0.05 (Kuehl, 2000). The statistical analysis was done using SAS software (SAS, Version 9.4, Cary, NC).

Multivariate statistics were performed using R (version 3.5.1; The R Foundation for Statistical Computing). Multivariate clustering analysis was done with the log_10_-transformed and Pareto scaling normalized data, with identified outlier observations removed. Missing values were estimated prior to multivariate analysis using k-nearest neighbor (KNN) in the MetaboAnalystR package (Chong and Xia, 2018). The total number of missing values within the inbred datasets was between 4.2% – 6.4% of all observations and between 4.8% – 6.4% for the hybrid datasets. Principal component analysis (PCA) was performed using the prcomp function (R stat package) for the inbred and hybrid experiments independently for each time point. A singular value decomposition data matrix was applied to each normalized dataset. When the unsupervised PCA identified a clear treatment separation, a supervised partial least square – discriminant analysis (PLS-DA) was performed, with the mixOmics package (Rohart *et al.*, 2017). The number of latent variables included in the model was selected by testing the predictability value (Q^2^) using an increasing number of latent variables from 1 to 10. The relative importance of the metabolites in the models was summarized using PLS-DA loadings, with significance considered when the contribution was greater than 0.1.

Pearson linear correlation analysis was done to investigate correlations between metabolite content and physiological traits (chlorophyll and LMA) based on the class separation observed in the multivariate clustering. Correlations with an adjusted *p* value (False Discovery rate, (Benjamini and Hochberg, 1995) of 0.05 or less, and a correlation coefficient of absolute value |0.55| or greater were considered significant and biologically meaningful in this analysis.

Statistical analyses of sterol:chlorophyll (relative concentration/ 100 mg DW: μg/cm^2^) and day of year (DOY) were performed using linear regression (Proc Reg; SAS). An F statistic was used to test if treatments had significantly different slopes in the regression, with each metabolite analyzed independently. Differences in slopes between ambient and elevated [O_3_] were considered significant when *p* < 0.05.

## Results

### Maize leaf metabolomic profiles in ambient and elevated [O_3_]

Metabolomic profiles of the leaf subtending the ear from two inbred lines (B73 and Mo17) and the hybrid cross (B73 x Mo17) were measured in ambient and elevated [O_3_] on three dates in 2015. These three time points captured the initial and middle stages of leaf senescence, based on leaf chlorophyll content (Fig. 1A-C). Both inbred lines and the hybrid showed acceleration of senescence, but it was more pronounced in the hybrid, which lost 25% more chlorophyll over time in elevated [O_3_] compared to ambient [O_3_] (Fig. 1C). Metabolite analysis detected 41, 45 and 46 annotated metabolites in the inbred lines at the three time points (Table S1), and 51, 49 and 56 metabolites at each time point in B73 x Mo17 (Table S2).

**Figure 1.**
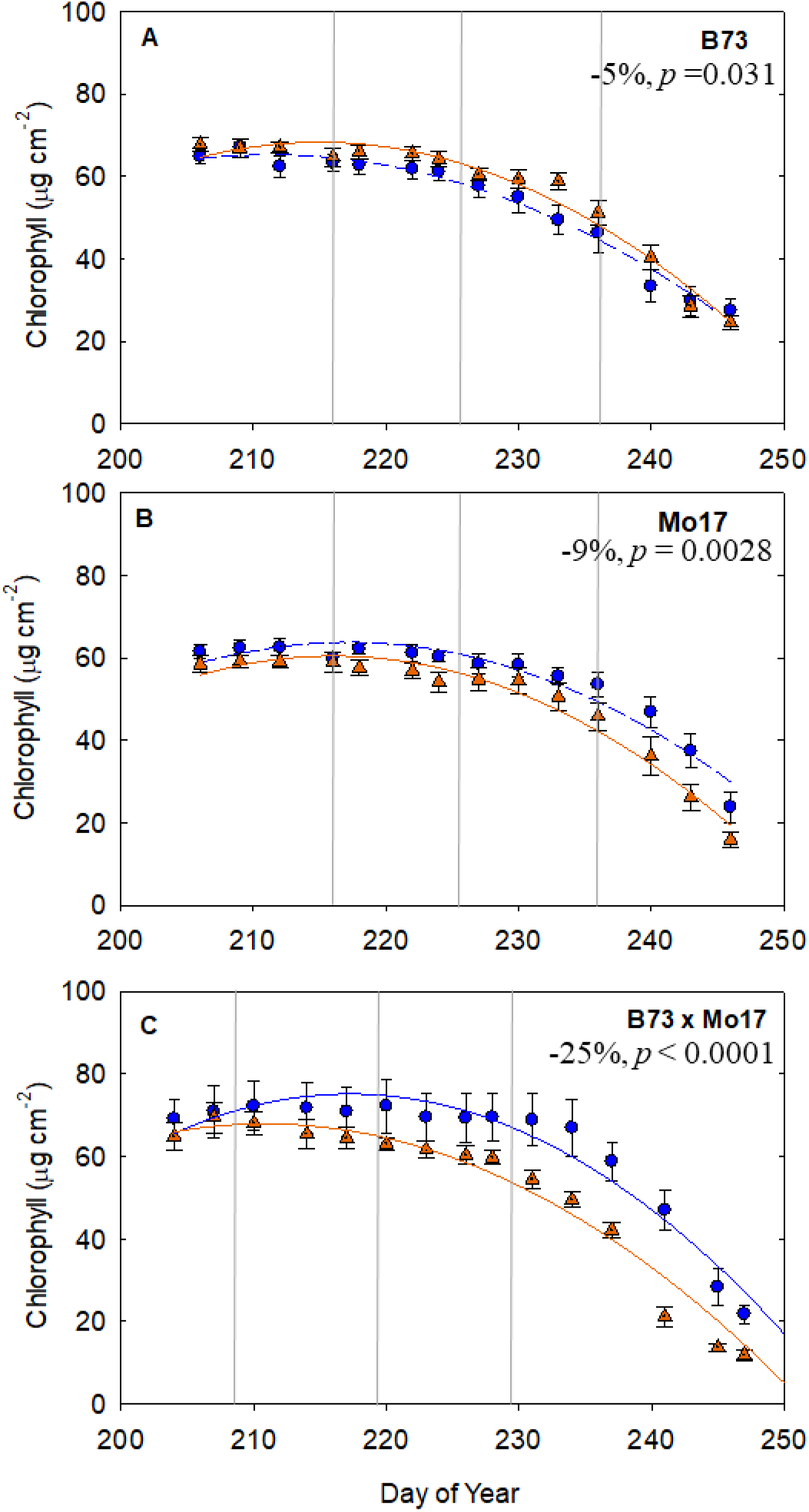
Chlorophyll content estimated from SPAD measurements of the leaf subtending the ear from leaf maturity through senescence for hybrid line B73 x Mo17 (A), and inbred lines B73 (B) and Mo17 (C) measured at ambient (blue symbols) and elevated [O_3_] (orange symbols). The area under the curve was integrated to calculate the percentage change in chlrophyll content over the lifetime of the leaf, and is shown in the top right of each plot, along with significance values. Vertical lines indicate the measurment dates for leaf metabolite content. Each point indicates the median value within each plot at each sampling date.

The leaf metabolite profile of the inbred lines did not show a significant response to elevated [O_3_] in any of the time points (Fig. 2A-C). This was true whether both inbred lines were analyzed together (Fig. 2A-C) or independently (data not shown). All time points showed a strong genotype separation between B73 and Mo17 along the first principle component. Similarly, statistical tests (ANOVA) identified 22, 37, and 24 metabolites in time point A, B, and C, with significant differences in content between genotypes (Table S3). In contrast, the content of only 3 metabolites differed in ambient and elevated [O_3_] (Table S3).

**Figure 2.**
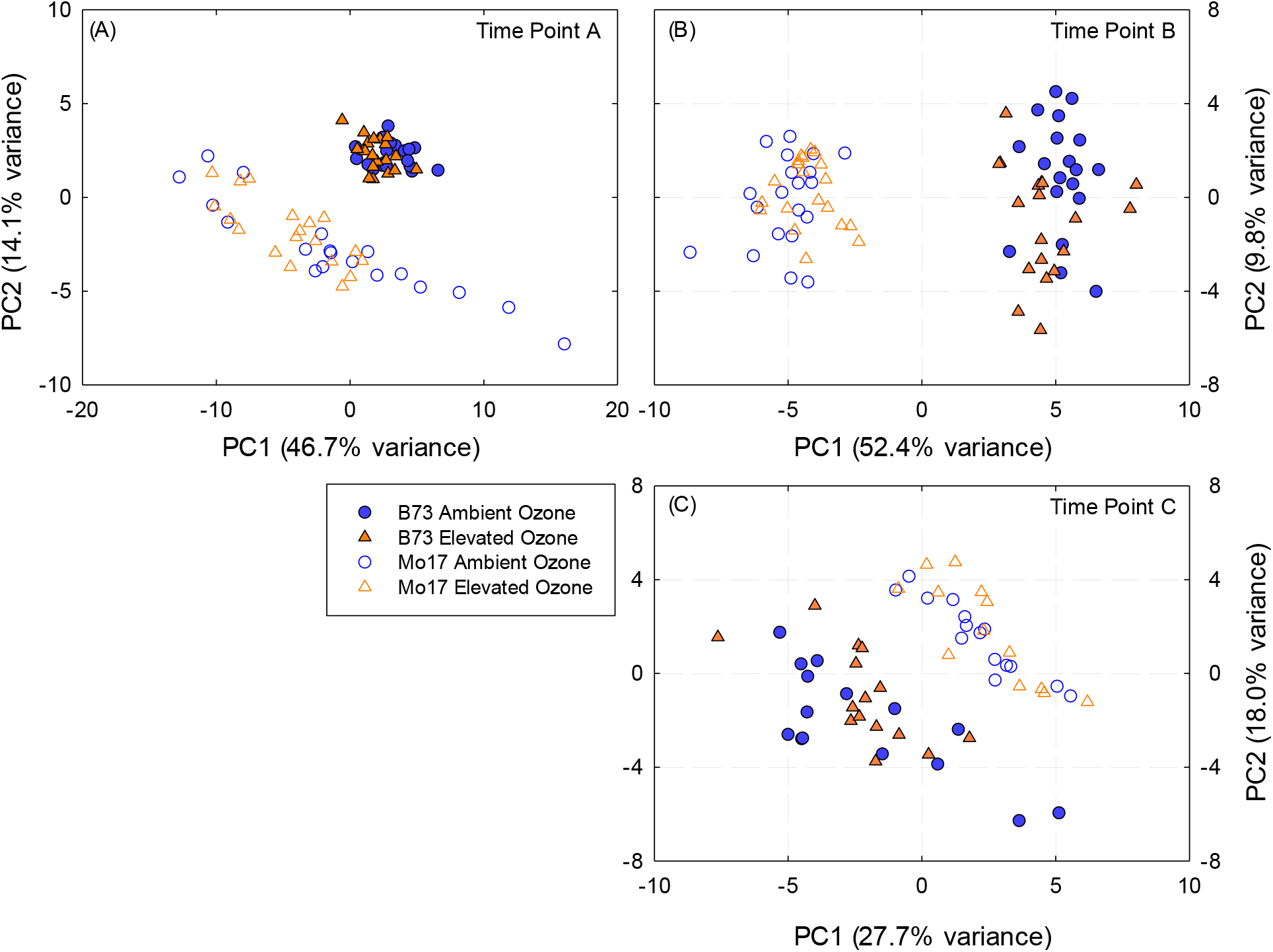
Principle component analysis plot of the metabolomic profile of inbred lines B73 (filled symbols) and Mo17 (open symbols) grown at ambient (blue symbols) and elevated [O_3_] (orange symbols) sampled at time points (A) DOY: 216, (B) DOY: 225, and (C) DOY: 236. n=20 for each ambient and elevated [O_3_] treatment and timepoint.

Multivariate clustering of the leaf metabolite profile of B73 x Mo17 across time points showed a clear shift in leaf metabolism under elevated [O_3_] as the leaves aged (Fig. 3). When the PCA plots revealed strong discrimination between ambient and elevated [O_3_], partial least square-discriminant analysis (PLS-DA) was performed to identify maximum separation among treatments and to rank individual metabolite contributions to separation of ambient and elevated [O_3_] (Figure S3). On the first date of sampling when the flag leaf was relatively young (time point A, DOY 208), the metabolite pools in B73 x Mo17 did not differ in ambient and elevated [O_3_] (Fig. 3A). As leaves aged in elevated [O_3_], both PCA and PLS-DA revealed differences in the metabolite profiles of leaves grown in ambient and elevated [O_3_] (Fig. 3B, 3C). Statistical tests (1-way ANOVA) identified 20 metabolites with significant differences in content between ambient and elevated [O_3_] in time point B, and 12 metabolites with significant differences in time point C (Table S4). Secondary metabolites quinic acid, stigmasterol, and α-tocopherol were greater in elevated [O_3_] in both time points (Fig. 4B-C), while sitosterol was greater in ambient [O_3_] (Fig. 4B-C). Fatty acids palmitate and linoleate were also greater in in ambient [O_3_] (Fig. 4E-F). The sugar, benzyl glucopyranoside, and two TCA cycle compounds, citric acid and malic acid, were greater in elevated [O_3_] in both time points. Pyruvate, which is converted to acetyl-CoA in the initial step of the citric acid cycle, was greater in ambient [O_3_] (Fig. 4E-F). Similarly, threonine, glycerol-3-P, and glycerophosphoglycerol were greater in ambient [O_3_] in time points B and C (Table S2).

**Figure 3.**
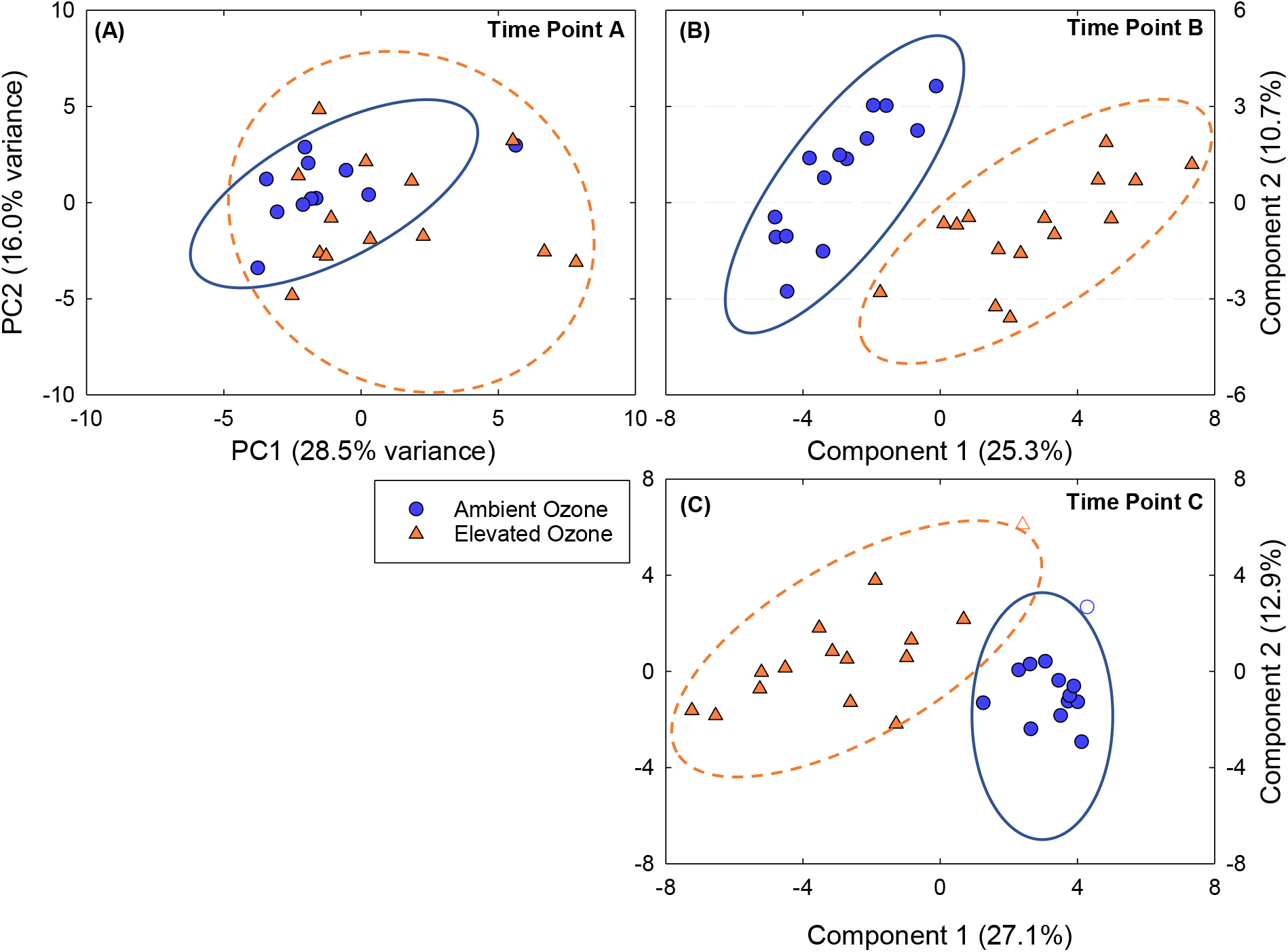
Multivariate clustering of the metabolomic profile of B73 x Mo17 hybrid at different time points. (A) Principle component analysis (PCA) of time point A (DOY: 208) showing no strong treatment separation; (B) Partial least square – discriminant analysis (PLS-DA) of time point B (DOY: 219), and (C) PLS-DA of time point C (DOY: 229). Ellipses show 95% confidence intervals and shapes without fill are outside the confidence ellipse for PLS-DA. For each time point ambient O_3_ n=15 and elevated O_3_ n=15.

**Figure 4.**
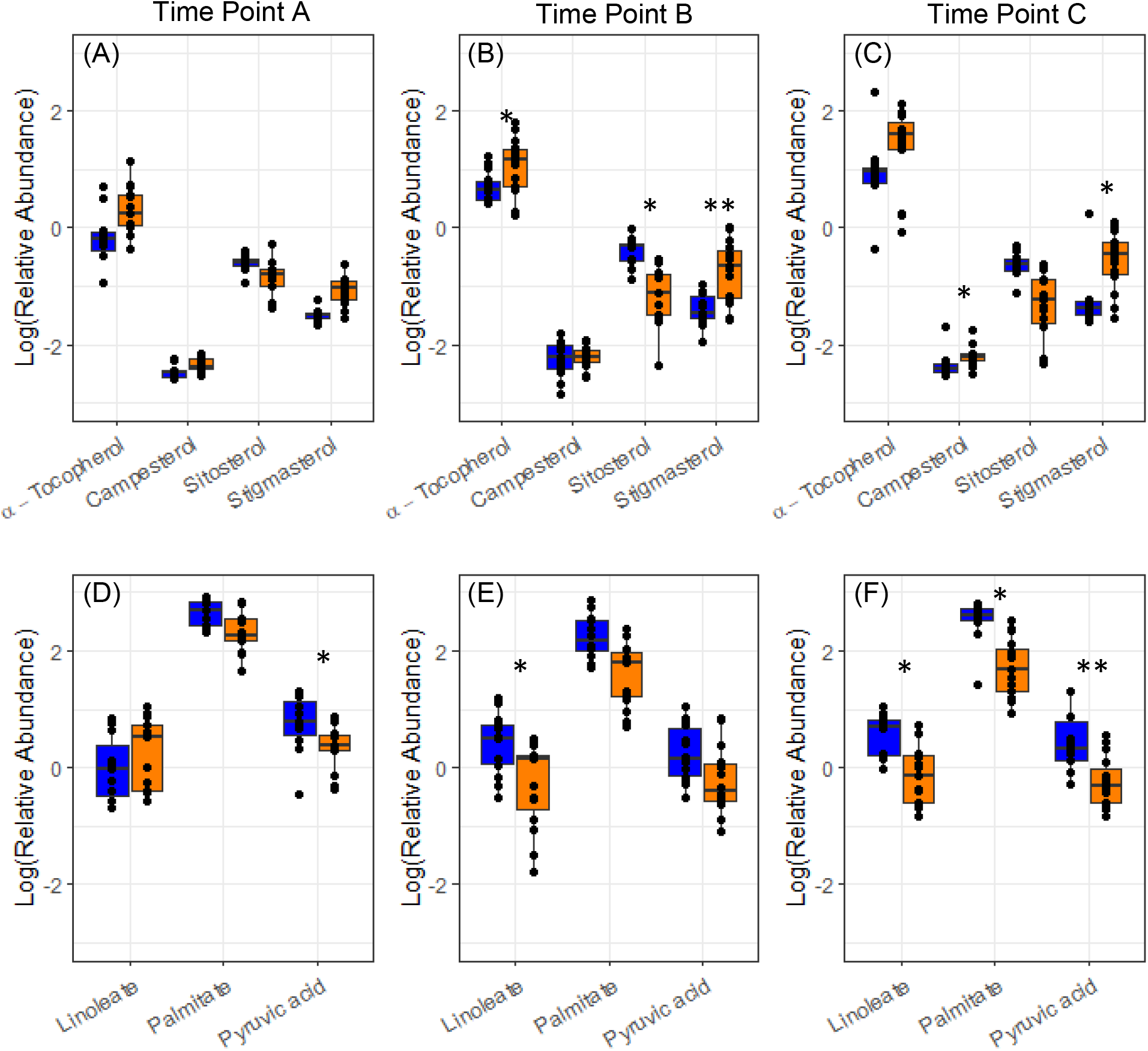
Leaf metabolite content for B73 x Mo17. Box plots show the relative abundance of metabolites sampled at time point A (A,D), time point B (B,E), and time point C (C,F). Statistical analyses for all metabolites are provided in Supplementary Table 3. Blue represents ambient [O_3_] and orange represents elevated [O_3_]. Black dots show individual values from each sample. (* p < 0.05, ** p<0.01).

The ratios of α-tocopherol, campesterol, and stigmasterol to chlorophyll, used as indices of senescence (Li *et al*., 2017), increased over time in B73 x Mo17, especially in elevated [O_3_] (Fig. 5A-C). These trends were due to both the decline in chlorophyll (Fig. 1) and an increase in α-tocopherol, campesterol, and stigmasterol content in aging leaves, especially in elevated [O_3_] (Fig. 4). A significant difference between ambient and elevated [O_3_] in the accumulation of α-tocopherol relative to chlorophyll (Fig. 5A) was observed, along with trends towards greater accumulation of stigmasterol and campesterol relative to chlorophyll (Fig. 5B, 5C). The reverse trend was seen in the sitosterol:chlorophyll ratio with decreasing ratios in the elevated O_3_ condition (Fig. 5D). The inbred lines showed an age-dependent increase in α-tocopherol:chlorophyll ratio, but no effect of elevated [O_3_] (Fig. 6).

**Figure 5.**
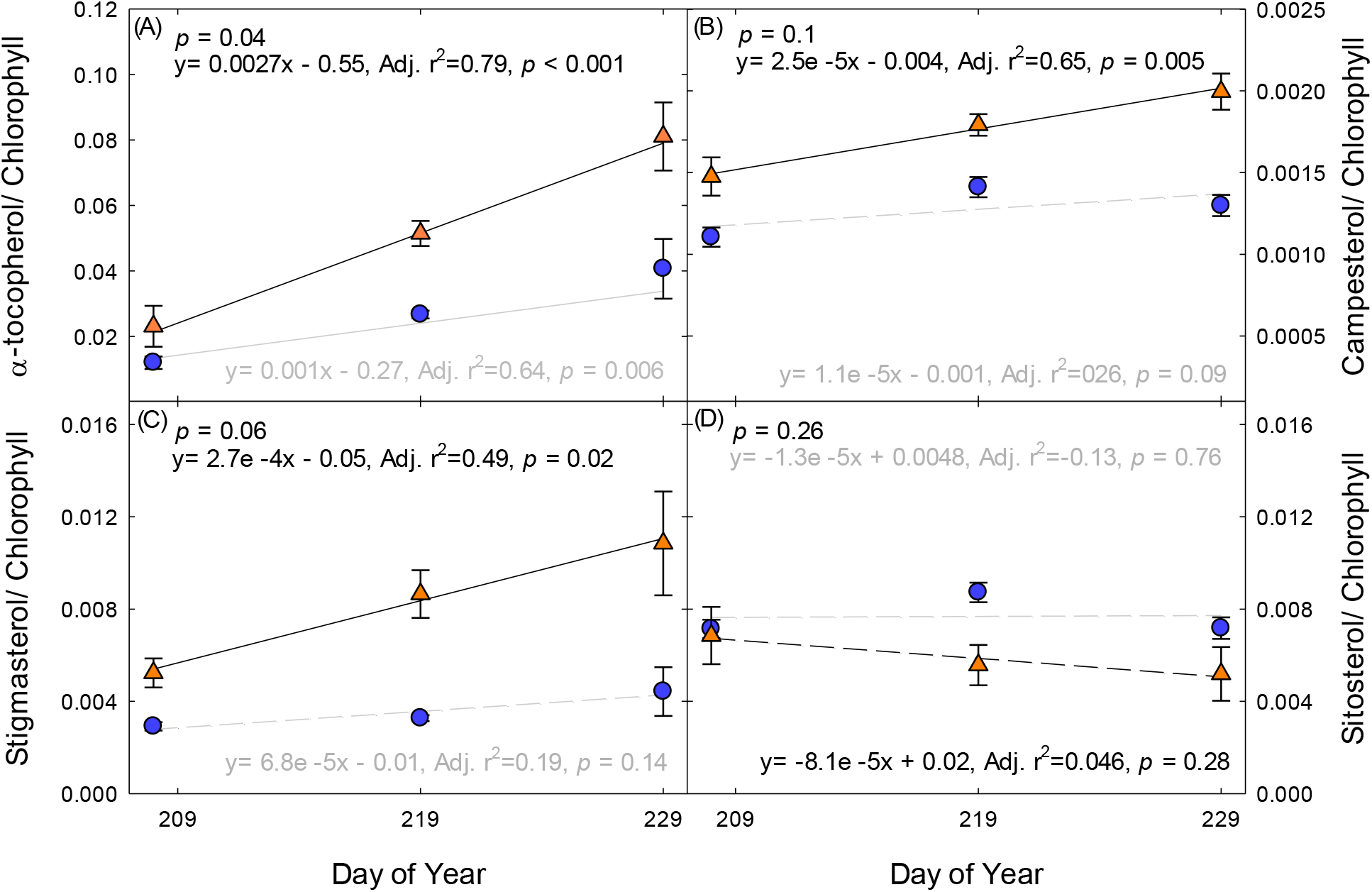
Ratio of leaf metabolite (relative concentration / 100 mg DW) content to chlorophyll content (μg/cm^2^) over time for hybrid B73 x Mo17. (A) α-tocopherol/chlorophyll ratio; (B) campesterol/chlorophyll ratio; (C) stigmasterol/chlorophyll ratio; (D) sitosterol/ chlorophyll ratio. Solid lines indicate statistically significant relationships, while dashed lines are not significant. *p*-value in the top left indicates significantly different slopes in the linear regressions in ambient (blue symbols, grey lines) and elevated [O_3_] (orange symbols, black lines). Each point indicates the median ratio of all ambient or elevated samples (n =15 per treatment).

**Figure 6.**
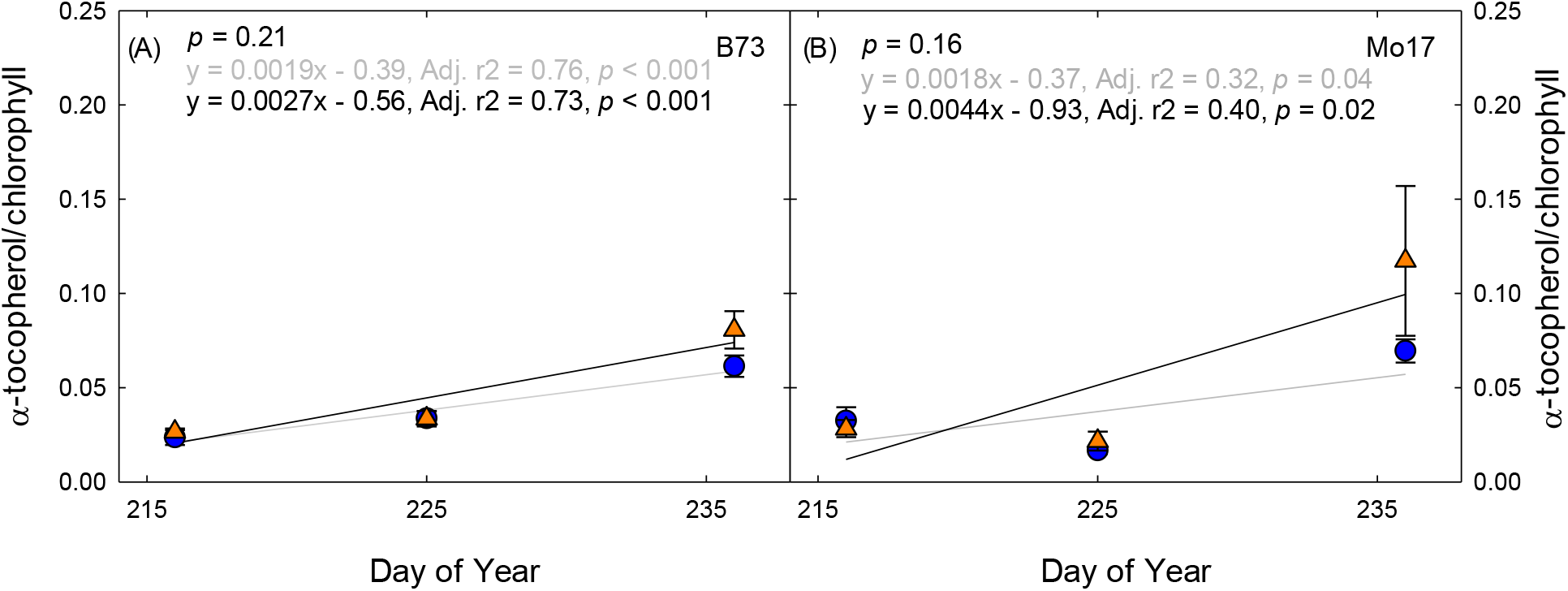
Ratio of α-tocopherol (relative concentration/ 100 mg DW) to chlorophyll (μg/cm^2^) content measured over time in (A) B73 and (B) Mo17 grown at ambient (blue symbols) and elevated [O_3_] (orange symbols). *p*-value in the top left indicates the test of differences in slope of the regression line between ambient and elevated [O_3_]. Each point indicates the median ratio of all ambient or elevated samples (n =20 per treatment).

### LMA and yield correlations with leaf metabolite content

LMA was not significantly affected by elevated [O_3_] in either inbred line at any time point, and was only significantly lower in elevated [O_3_] in the hybrid at the last sampling time (Table 1). Seed yield was not significantly affected by elevated [O_3_] in either inbred line, but was nearly 30% lower in elevated [O_3_] in the hybrid (Table 1).

**Table 1.**
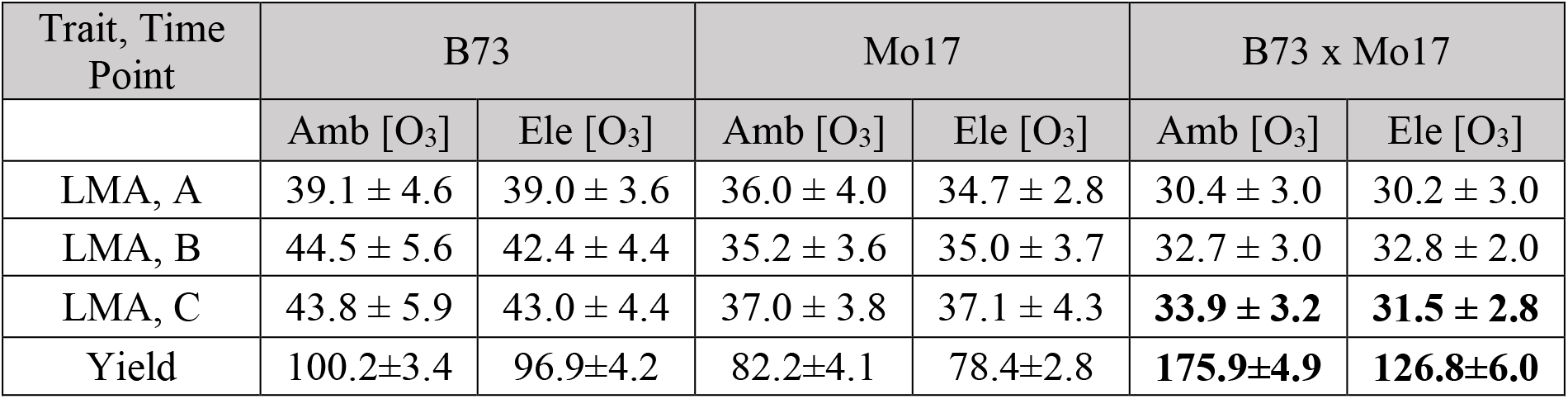
Leaf mass per unit area (LMA, g m^−2^) measured at three time points (A, B, C) on the leaf subtending the ear. Inbred leaves were sampled on 4 Aug (A), 13 Aug (B) and 24 Aug (C). Hybrids were sampled on 27 July (A), 7 Aug (B), and 17 Aug (C). Yield (kernel mass per plant, g plant^−1^) was measured at maturity in ambient (Amb) and elevated [O_3_] (Ele). Bold font indicates significant differences in ambient and elevated [O_3_] (*p*< 0.05).

Correlations between metabolite content and LMA or yield were tested separately for inbred and hybrid maize lines. In the inbred lines, itaconic acid measured at time point A was positively correlated with yield (Table S5). Stigmasterol and LMA were negatively correlated at time point B in B73, while raffinose and LMA showed a positive correlation at time point C. The strongest relationships within Mo17 were between malic acid and LMA in time point A, and sucrose and LMA in time point C (Table S5).

In B73 x Mo17, there was no treatment separation between ambient and elevated [O_3_] at time point A, so correlations were done across O_3_ treatments (Table S6). Stigmasterol, campesterol, and ethanolamine contents measured in recently mature leaves (time point A) were negatively correlated with yield, while alanine was positively correlated with yield (Table 2). Multivariate clustering for time points B and C showed significant differences in ambient and elevated [O_3_] in B73 x Mo17, so correlations were done separately for each O_3_ treatment (Table 3). Benzyl glucopyranoside, campesterol, glycerohexose, and malic acid content were negatively correlated with yield in elevated [O_3_], but not in ambient [O_3_] (Table 3). Scyllo inositol was positively correlated with yield in ambient [O_3_], and negatively correlated with yield in elevated [O_3_] (Table 3). Ferulic acid, itaconic acid, and citric acid measured at time point C (when leaves were senescing) were positively correlated with yield in ambient [O_3_], but not in elevated [O_3_] (Table 4).

**Table 2.**
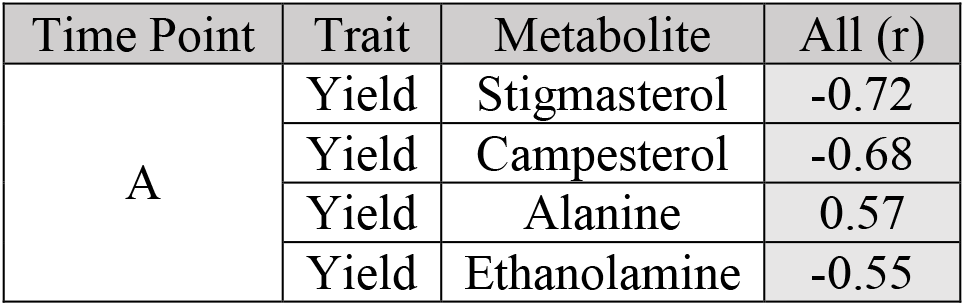
Significant Pearson linear correlations (*p*<0.05) between yield and metabolite content for hybrid line, B73 x Mo17. Linear correlation statistics for time point A (DOY: 208) contain all samples from both ambient and elevated [O_3_].

**Table 3.** Significant Pearson linear correlations (*p*<0.05) of yield and leaf mass per unit area (LMA) with metabolite content sampled at time point B (DOY: 219) for hybrid line, B73 x Mo17 in ambient and elevated [O_3_].

| Time Point | Trait | Metabolite | Ambient (r) | Elevated (r) |
| --- | --- | --- | --- | --- |
| B | Yield | Benzyl glucopyranoside | 0.15 | -0.60 |
|  | Yield | Campesterol | 0.01 | -0.58 |
|  | Yield | Glycerohexose | -0.26 | -0.56 |
|  | Yield | Malic acid | 0.14 | -0.55 |
|  | Yield | Inositol, scyllo | 0.56 | -0.54 |
| B | LMA | Sitosterol | -0.58 | 0.55 |
|  | LMA | Threonic acid | -0.50 | -0.57 |
|  | LMA | Benzyl glucopyranoside | -0.57 | -0.32 |

**Table 4.** Significant Pearson linear correlations (*p*<0.05) of yield and leaf mass per unit area (LMA) with metabolite content sampled at time point C (DOY: 229) for hybrid line, B73 x Mo17 in ambient and elevated [O_3_].

| Time Point | Trait | Metabolite | Ambient (r) | Elevated (r) |
| --- | --- | --- | --- | --- |
| C | Yield | Ferulic acid | 0.81 | 0.02 |
|  | Yield | Itaconic acid | 0.68 | -0.04 |
|  | Yield | Citric acid | 0.57 | -0.29 |
| C | LMA | α-tocopherol | 0.14 | 0.60 |
|  | LMA | Maltose | -0.25 | 0.60 |
|  | LMA | Malonic acid | -0.59 | 0.05 |
|  | LMA | o-coumaric acid | -0.67 | 0.03 |
|  | LMA | Itaconic acid | -0.05 | -0.64 |
|  | LMA | Pyruvic acid | 0.21 | -0.63 |
|  | LMA | Ribose | 0.11 | -0.60 |
|  | LMA | Alanine | -0.12 | -0.58 |
|  | LMA | Pyroglutamic acid | -0.40 | -0.57 |

Sitosterol and LMA showed opposite correlations in ambient and elevated [O_3_] when measured at time point B (Table 3) for B73 x Mo17. Benzyl glucopyranoside and LMA were negatively correlated in time point B in ambient [O_3_] (Table 3). In time point C, LMA and α-tocopherol were positively correlated in elevated [O_3_], but not in ambient [O_3_] (Table 4). Two significant linear correlations were identified in the ambient [O_3_] treatment with LMA, o-coumaric acid and malonic acid (Table 4). Itaconic acid, pyruvic acid, ribose, alanine, and pyroglutamic acid were all negatively correlated with LMA in time point C in elevated [O_3_] (Table 4).

## Discussion

Field metabolomics can be a powerful approach for profiling the metabolite changes of plants in response to environmental stress (Wedow *et al.*, 2019; Melandri *et al.*, 2020). The physiological response of crops to O_3_ pollution has been well-studied, with elevated [O_3_] decreasing carbon assimilation, accelerating senescence and cell death, and reducing economic yield (Ainsworth, 2017). However, the metabolite profile underpinning the response to O_3_ in maize has not been investigated and could provide information for targets to improve O_3_ tolerance in maize and C_4_ crops. We found that maize metabolomics signatures were altered by elevated [O_3_] in a hybrid line (Fig. 2), but not in two inbred lines (Fig. 3). As predicted, the differences in metabolite signatures between ambient and elevated [O_3_] in the hybrid line emerged as leaves aged (Fig. 2). Notably, there was a significant increase in α-tocopherol and phytosterol content in elevated [O_3_], supporting the prediction that metabolites that quench oxidative stress and stabilize membranes are key biochemical responses to O_3_ stress.

The metabolomic profiles of inbred and hybrid leaves reflected differences in acceleration of senescence under elevated [O_3_]. In inbred lines and when chlorophyll content was similar in ambient and elevated [O_3_] in B73 x Mo17, the metabolomic profile was not different in ambient and elevated [O_3_] (Fig. 3, Fig. 2A). However, as chlorophyll was lost more rapidly at elevated [O_3_] during senescence in the hybrid, a clear separation in metabolite profiles between ambient and elevated [O_3_] was apparent (Fig. 2B-C). The inbred genotypes had no discernible difference in their metabolomic profiles in ambient and elevated [O_3_] at any of the time points, although B73 was very different from Mo17 (Fig. 3). These results agree with our prediction that a metabolomic signature develops with the accumulation of O_3_ damage in aging leaves and reflects altered biochemistry. Ozone-induced acceleration of senescence has been previously reported to impact grain yield losses due to a decrease in photosynthetic capacity and shortened leaf lifespan (Emberson *et al.*, 2018). A reduction in photosynthetic carbon assimilation in maize ear leaves during grain filling was greater in hybrid maize compared to inbred lines (Yendrek *et al.*, 2017a). Additionally, seed yield loss under elevated [O_3_] was 30.4% for B73 x Mo17, and only 5.0% for Mo17, and 2.1% for B73. Thus, the hybrid maize line showed greater response to elevated [O_3_] from the biochemical to the agronomic scale. This is despite the observation that stomatal conductance was not significantly altered by elevated [O_3_] in either B73 x Mo17 or the two inbred lines (Yendrek *et al*., 2017a), suggesting that differential stomatal sensitivity to O_3_ was not a primary driver of differences between the hybrid and inbred lines.

The metabolites contributing to the O_3_ treatment differences during leaf senescence in B73 x Mo17 included TCA cycle intermediates, and specialized metabolites such as phytosterols and fatty acids (Fig. 4). In the latter two time points, citric acid and malic acid contents were greater in elevated [O_3_]. This trend is consistent with increased flux through the TCA cycle and greater rates of mitochondrial respiration in plants exposed to O_3_ stress (Betzelberger *et al.*, 2012; Dizengremel *et al.*, 2012; Yendrek *et al.*, 2015). The shift in metabolism from photosynthetic carbon assimilation to the TCA cycle and mitochondrial respiration supplies energy for repair and detoxification of the plant cells against oxidative damage (Dizengremel, 2001). The additional influence of specialized secondary metabolites including α-tocopherol, benzyl glucopyranoside, stigmasterol, and quinic acid is consistent with a shift to defense and repair under elevated O_3_ stress (Fig. 4). Steryl esters (SEs), palmitate and linoleate, which are commonly found to function as membrane lipids were greater in ambient [O_3_], possibly reflecting differences in membrane fluidity or other membrane properties between ambient and elevated [O_3_].

We found that α-tocopherol increased to a greater extent as leaves aged in elevated [O_3_] in the hybrid B73 x Mo17 (Fig. 5A-C), but not in either inbred line (Fig. 6). The protective function of tocopherols to preserve cell membrane integrity during the final stages of leaf development has been well documented (Fryer, 1992; Falk and Munne-Bosch, 2010; Lira *et al.*, 2017). Leaf α-tocopherol content increases in response to abiotic stresses, such as high light (Krieger-Liszkay and Trebst, 2006; Lizarazo *et al*., 2010), drought (Munne-Bosch and Alegre, 2000; Ahkami *et al.*, 2019), and high temperature stress (Spicher *et al.*, 2017). In Mediterranean species, α-tocopherol showed greater environmental sensitivity to drought and high temperature stress compared to small antioxidant molecules (ascorbate, glutathione) and xanthophyll-cycle pigments (Fernández-Marín *et al*., 2017). Within the chloroplast, where highly reactive singlet oxygen (^1^O_2_) is formed, α-tocopherol is an essential scavenger protecting the thylakoid membranes against lipid peroxidation (Piller *et al.*, 2014). In high light conditions, the rapid turnover of α-tocopherol and plastoquinone have been correlated with the increased turnover rate of the D1 protein within PSII, thereby protecting the photosynthetic process (Krieger-Liszkay and Trebst, 2006). Experimental studies exposing plants to oxidative stresses have reported conflicting results regarding α-tocopherol levels in leaf extracts; beech leaves in full sun showed a significant increase over shade leaves, along with an acceleration in leaf senescence (García-Plazaola and Becerril, 2001). Snap bean showed a significant increase in α-tocopherol concentrations following exposure to elevated [O_3_]; however, the total concentration of α-tocopherol was not correlated with differences in O_3_ sensitivity between cultivars (Burkey *et al*., 2001). In contrast, spinach leaves exposed to elevated [O_3_] were observed to have a decrease or no change in α-tocopherol in leaf extracts (Calatayud *et al.*, 2003, 2004). In this study, the response of leaf α-tocopherol content to elevated [O_3_] varied with maize genotype and leaf age (Fig. 5A; Fig. 6). The conflicting results from previous studies regarding effects of elevated [O_3_] on leaf α-tocopherol content may be attributed to the timing of sampling and the progression of leaf senescence. We would not have identified a relationship between α-tocopherol and O_3_ in B73 x Mo17 had samples only been taking at the initial time point (Fig. 4A), but clearly the content increased as leaves aged in elevated [O_3_]. Elevated α-tocopherol concentrations in leaves exposed to elevated [O_3_] are potentially quenching ROS that form during the disassembly of the photosynthetic membranes during senescence and/or inhibiting lipid peroxidation (Rogers and Munne-Bosch, 2016). Manipulation of α-tocopherol content in leaves has been proposed as an efficient trait to improve dynamic responses to abiotic stress (Lizarazo *et al*., 2010). It would be interesting to test if transgenic approaches to increase α-tocopherol in aging leaves would specifically improve O_3_ tolerance.

Senescing hybrid leaves exposed to elevated [O_3_] showed a trend towards accumulating campesterol and stigmasterol (Fig. 5B, C) at the expense of sitosterol (Fig. 5D). Phytosterols are proposed to modulate membrane integrity to improve abiotic stress tolerance (Dufourc, 2008; Kuczynska *et al.*, 2019). In stress conditions stigmasterol and sitosterol interact with phospholipids to maintain the permeability and fluidity of the plasma membrane (Dalal *et al.*, 2016; Griebel and Zeier, 2010). Heat stressed hard fescues up-regulated ethyl sterol content, including stigmasterol and sitosterol, and the more heat tolerant variety showed a significantly greater increase in stigmasterol compared to a heat sensitive variety (Wang *et al.*, 2017). In the current study, we found greater concentrations of stigmasterol and campesterol in hybrid leaves exposed to elevated [O_3_], but not in inbred lines (Fig. 5; Supplementary Table 5,6). Genotypic variation in phytosterol concentrations has been documented in wheat (Nurmi *et al.*, 2010), rice (Kumar *et al.*, 2018), and potato (Hancock *et al.*, 2014), and genetic variation in the concentrations of phytosterols during drought was associated with differences in stress tolerance (Kumar *et al.*, 2015). It is interesting that in elevated [O_3_], increased phytosterol content only occurred in the hybrid genotype that also showed accelerated senescence and significant yield loss, not in the inbred lines. This suggests that increased phytosterol content was not associated with O_3_ tolerance, instead with changes to membranes in aging leaves. Furthermore, campesterol is a precursor of brassinosteroids (Fujioka and Yokota, 2003), and brassinosteroid signaling mutants display a delayed senescence phenotype (Clouse and Sasse, 1998). Thus, the accumulation of campesterol in aging leaves exposed to elevated [O_3_] in B73 x Mo17 could further promote leaf senescence. Such fluctuations in the levels of one phytosterol at the expense of another are proposed to alter plant responses to environmental stimuli (Aboobucker and Suza, 2019). Our work also supports the hypothesis that the balance of various phytosterols may be a key signaling response for additional cellular defenses (Griebel and Zeier, 2010; Schaller, 2003).

Previous experiments have identified metabolites associated with maize grain yield under drought and heat stress (Obata *et al*., 2015), and here we investigated potential metabolite markers associated with yield under elevated [O_3_] stress. In the B73 x Mo17 genotype, stigmasterol and campesterol showed negative correlations to yield at the first time point, before any treatment effect could be identified (Table 2). In the second time point, campesterol content was negatively correlated with yield in the elevated [O_3_] treatment, but not in ambient [O_3_] (Table 3). There were no metabolites associated with yield in both ambient and elevated [O_3_] in either time point B or C (Table 3; Table 4), when PCA and PLS-DA analysis revealed strong treatment effects on the leaf metabolite profile. This supports previous work that identified different metabolite correlations with yield under drought stress compared to heat stress (Obata *et al*., 2015). LMA and α-tocopherol were positively correlated under elevated [O_3_] (Table 4). A reduction in LMA is generally linked to decreasing leaf nitrogen content and leaf area under elevated O_3_ conditions (Oikawa and Ainsworth, 2016). The positive relationship identified suggests a potential protective role of α-tocopherol in stabilizing thylakoid membranes and preventing excessive degradation of chlorophyll under elevated [O_3_]. The strong relationships identified in this study provide potential metabolic signatures for improving O_3_ tolerance, and provide metabolite markers that can now be screened across a diverse genetic background.

Field experiments can be more variable than controlled environment experiments (Lovell *et al.*, 2016), and metabolites respond rapidly to environmental changes (Caldana *et al.*, 2011), both of which present challenges for field metabolomic studies. Recent criticism of metabolomics data includes the low repeatability and strong effects of environmental perturbations on plant metabolism (Tucker *et al.*, 2020). We found that environmental conditions influenced the metabolomic profiles; yet there were clear patterns of response in hybrid maize. Moreover, we discovered that maize inbred lines have significantly different metabolite profiles from one another, but lacked a response to elevated [O_3_]. Our study emphasized that metabolomic profiling is a vital tool that can be used alongside additional ‘omic’ and standard physiological measurements to improve understanding of plant metabolic responses to stress. Despite the dynamic and variable nature of field-based metabolomics, we identified novel markers of O_3_ response in hybrid maize, which can be broadly tested across diverse germplasm.

## Acknowledgments

This work was supported by a grant from the NSF Plant Genome Research Program (PGR-1238030). We thank Lauren McIntyre and Pat Brown for help with the experimental field design, Kannan Puthuval, Brad Dalsing, and Chad Lance for management of the FACE rings and fumigation, and Craig Yendrek, Chris Montes, Nicole Choquette, Taylor Pederson, Pauline Lemonnier, Crystal Sorgini, Alvaro Sanz-Saez, John Regan, Ben Thompson, Matthew Kendzior, and Mark Lewis for help with sampling. Any opinions, findings and conclusions or recommendations expressed in this publication are those of the author(s) and do not necessarily reflect the views of the U.S. Department of Agriculture. Mention of trade names or commercial products in this publication is solely for the purpose of providing specific information and does not imply recommendation or endorsement by the U.S. Department of Agriculture. USDA is an equal opportunity provider and employer.

## Author Contribution

JMW and EAA conceptualized the study. JMW performed metabolomic and statistical analysis. CHB collected and analyzed leaf chlorophyll data with EAA. LRA and ADBL collected yield data. ADBL and EAA designed the field experiment. JMW and EAA wrote the manuscript with input from all authors.

## Figure legends

**Figure 1. Chlorophyll content in maize leaves.** Chlorophyll content was estimated from SPAD measurements of the leaf subtending the ear from leaf maturity through senescence for hybrid line B73 x Mo17 (A), and inbred lines B73 (B) and Mo17 (C) measured at ambient (blue symbols) and elevated [O_3_] (orange symbols). The area under the curve was integrated to calculate the percentage change in chlrophyll content over the lifetime of the leaf, and is shown in the top right of each plot, along with significance values. Vertical lines indicate the measurment dates for leaf metabolite content. Each point indicates the median value within each plot at each sampling date.

**Figure 2.** Principle component analysis plot of the metabolomic profile of inbred lines B73 (filled symbols) and Mo17 (open symbols) grown at ambient (blue symbols) and elevated [O_3_] (orange symbols) sampled at time points (A) DOY: 216, (B) DOY: 225, and (C) DOY: 236. For each timepoint and treatment, n=20.

**Figure 3.** Multivariate clustering of the metabolomic profile of B73 x Mo17 hybrid at different time points. (A) Principle component analysis (PCA) of time point A (DOY: 208) showing no strong treatment separation; (B) Partial least square – discriminant analysis (PLS-DA) of time point B (DOY: 219), and (C) PLS-DA of time point C (DOY: 229). Ellipses show 95% confidence intervals and shapes without fill are outside the confidence ellipse for PLS-DA. For each time point ambient O_3_ n=15 and elevated O_3_ n=15.

**Figure 4.** Leaf metabolite content for B73 x Mo17. Box plots show the relative abundance of selected metabolites sampled at time point A (A,D), time point B (B,E), and time point C (C,F). Statistical analyses for all metabolites are provided in Supplementary Table 3. Blue represents ambient [O_3_] and orange represents elevated [O_3_]. Black dots show individual values from each sample. (* p < 0.05, ** p<0.01).

**Figure 5.** Ratio of leaf metabolite (relative concentration / 100 mg DW) content to chlorophyll content (μg/cm^2^) over time for hybrid B73 x Mo17. (A) α-tocopherol/chlorophyll ratio; (B) campesterol/chlorophyll ratio; (C) stigmasterol/chlorophyll ratio; (D) sitosterol/ chlorophyll ratio. Solid lines indicate statistically significant relationships, while dashed lines are not significant. *p*-value in the top left indicates significantly different slopes in the linear regressions in ambient (blue symbols, grey lines) and elevated [O_3_] (orange symbols, black lines). Each point indicates the median ratio of all ambient or elevated samples (n =15 per treatment).

**Figure 6.** Ratio of α-tocopherol (relative concentration/ 100 mg DW) to chlorophyll (μg/cm^2^) content measured over time in (A) B73 and (B) Mo17 grown at ambient (blue symbols) and elevated [O_3_] (orange symbols). *p*-value in the top left indicates the test of differences in slope of the regression line between ambient and elevated [O_3_]. Each point indicates the median ratio of all ambient or elevated samples (n =20 per treatment).

## Supplemental Figures & Tables

**Figure S1:** Google Earth image of the SoyFACE field site showing the inbred and hybrid plots. The experimental design for one inbred and one hybrid plot is also shown. Different colors represent the 5 sectors of the ring. Each genotype was sampled from all 5 sectors of the ring for metabolite analysis.

**Figure S2:** Experimental analysis outline. Blue shaded boxes show pipeline of hybrid metabolomic profiling. Green shaded boxes show pipeline of inbred metabolomic profiling. Peach shaded boxes show pipeline for linear correlations of metabolites and traits, hybrid and inbred datasets were again analyzed independently along with time points. For the LMA and seed yield ANOVA analysis, the linear model is equivalent to what is shown for the metabolomic profiling but not shown in the diagram. Terms for the linear model are as follows: treatment (т), blocks (B), genotype (G), error (е).

**Figure S3:** PLS-DA loading plot for specific metabolites sampled from B73 x Mo17 for the first component for (A) time point B (DOY 219) and (B) time point C (DOY 229). Bars indicate the expression value for each metabolite in blue (ambient [O_3_]) and orange (elevated [O_3_]). Underlined metabolites are commonly expressed within an [O_3_] treatment across the two sampling dates. The numbers indicate general metabolite classification: (1) alcohol, (2) amino acid, (3) fatty acid, (4) organic acid (5) polyol, (6) sugar, (7) specialized metabolite (8) other.

**Table S1:** Mean, standard error, minimum and maximum metabolite content measured in ambient and elevated [O_3_] in the hybrid B73 x Mo17 at timepoint A (DOY 208), timepoint B (DOY 219) and timepoint C (DOY 229).

**Table S2:** Mean, standard error, minimum and maximum metabolite content measured in inbred lines B73 and Mo17 at timepoint A (DOY 216), timepoint B (DOY 225) and timepoint C (DOY 236).

**Table S3:** Statistical analysis (ANOVA) for individual metabolites, LMA and yield for the inbred lines B73 and Mo17.

**Table S4:** Statistical analysis (ANOVA) for individual metabolites, LMA and yield from the hybrid line B73 x Mo17.

**Table S5:** Linear correlations between grain yield or LMA and metabolite content in inbred lines B73 and Mo17 (grey).

**Table S6:** Linear correlations between grain yield or LMA and metabolite content in hybrid line B73 x Mo17.

## Notes

### Competing Interest Statement

The authors have declared no competing interest.

