## Supplemental Figures for "Age-dependent increase in α-tocopherol and phytosterols in maize leaves exposed to elevated ozone pollution"

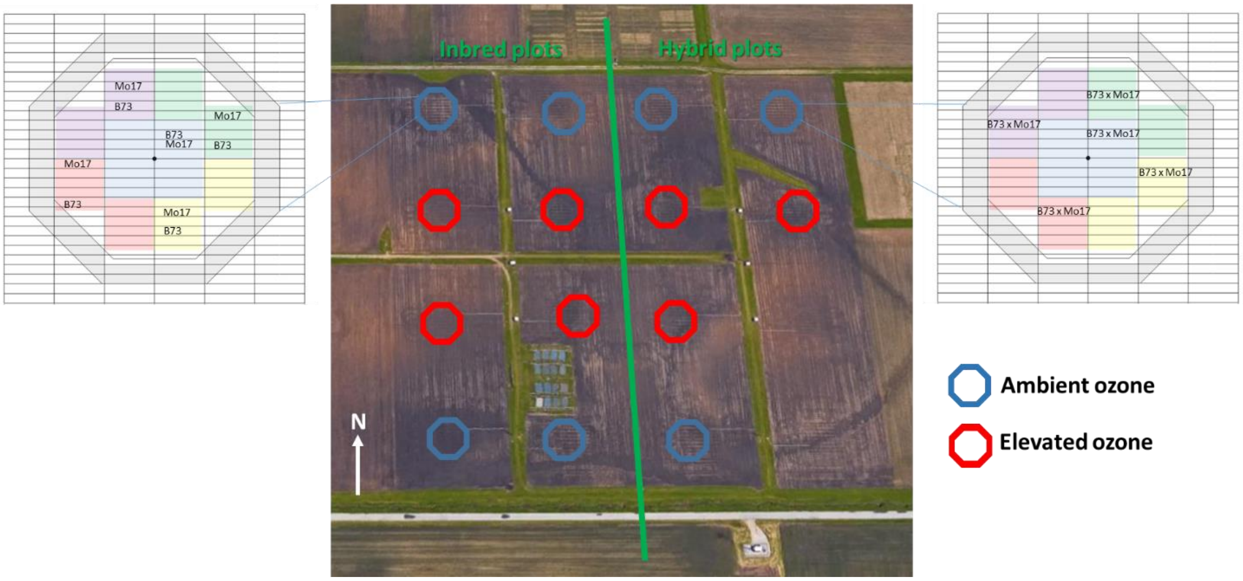


**Figure S1:** Google Earth image of the SoyFACE field site showing the inbred and hybrid plots. The experimental design for one inbred and one hybrid plot is also shown. Different colors represent the 5 sectors of the ring. Each genotype was sampled from all 5 sectors of the ring for metabolite analysis.

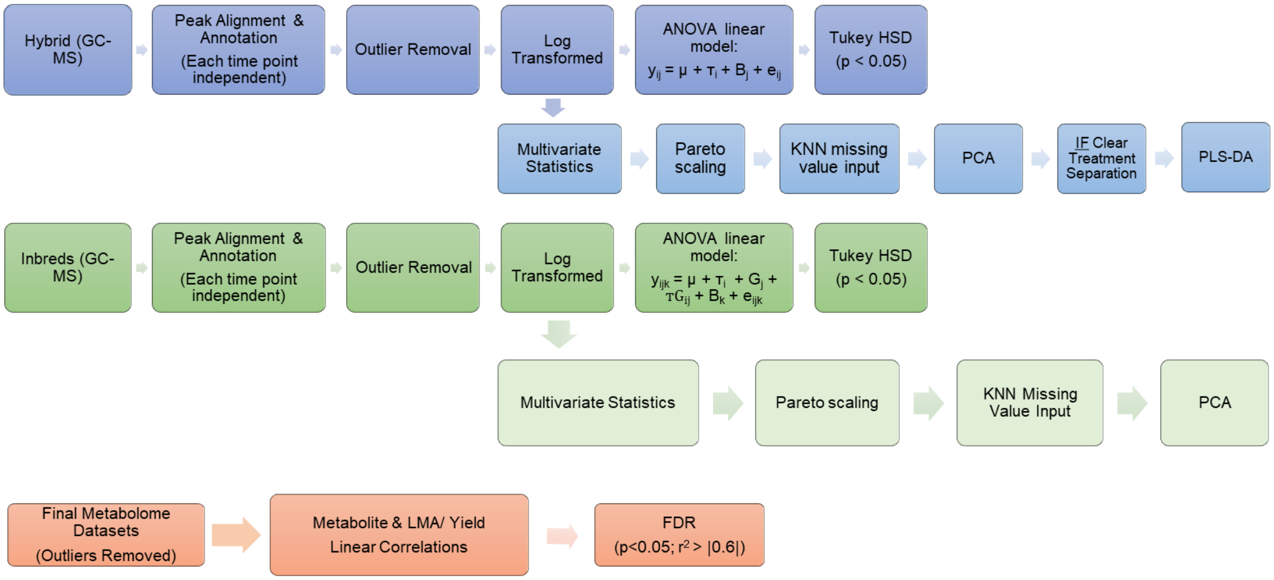


**Figure S2:** Experimental analysis outline. Blue shaded boxes show pipeline of hybrid metabolomic profiling. Green shaded boxes show pipeline of inbred metabolomic profiling. Peach shaded boxes show pipeline for linear correlations of metabolites and traits, hybrid and inbred datasets were again analyzed independently along with time points. For the LMA and seed yield ANOVA analysis, the linear model is equivalent to what is shown for the metabolomic profiling but not shown in the diagram. Terms for the linear model are as follows: treatment (τ), blocks (B), genotype (G), error (e).

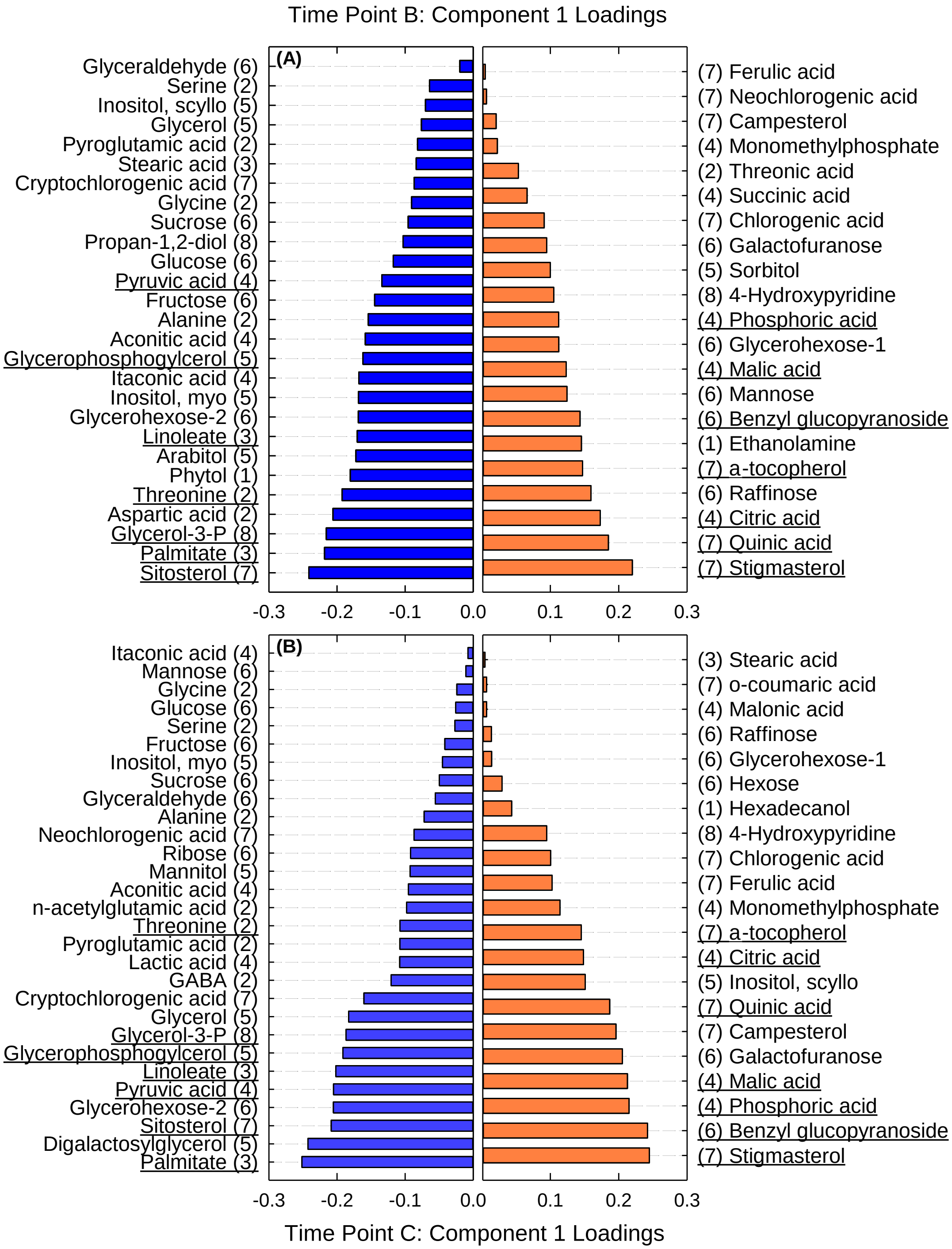


**Figure S3:** PLS-DA loading plot for specific metabolites sampled from B73 x Mo17 for the first component for (A) time point B (DOY 219) and (B) time point C (DOY 229). Bars indicate the expression value for each metabolite in blue (ambient [$O_3$]) and orange (elevated [$O_3$]). Underlined metabolites are commonly expressed within an [$O_3$] treatment across the two sampling dates. The numbers indicate general metabolite classification: (1) alcohol, (2) amino acid, (3) fatty acid, (4) organic acid (5) polyol, (6) sugar, (7) specialized metabolite (8) other.
